# Integrin α6β4 is downregulated in mutant IDH1 oligodendrogliomas, promotes glioma growth, and associates with a worse outcome in glioma patients

**DOI:** 10.1101/726489

**Authors:** Rachel L. Stewart, Min Chen, Erin E. Mulkearns-Hubert, Justin Lathia, Kathleen L. O’Connor, Craig Horbinski

## Abstract

The integrin α6β4 is a laminin receptor that associates with aggressive behavior in many cancers, though its significance in gliomas has not been established. Mutations in *isocitrate dehydrogenase 1* and *2* (IDH^mut^) are common in gliomas, especially 1p/19q co-deleted oligodendrogliomas, and correlate with improved patient survival. We sought to determine whether there is a link between integrin β4 and malignant behavior in gliomas. An inverse relationship was identified between IDH^mut^ and integrin β4 expression in a glioma TMA (*P* < 0.0001) and in TCGA grade II-IV gliomas (*P* < 0.0001). Methylation of CpG sites in the *ITGB4* promoter is significantly higher in IDH^mut^ gliomas when compared to wild-type tumors, and both *ITGB4* methylation and reduced *ITGB4* mRNA are not only most prominent in the 1p/19q co-deleted subset of IDH^mut^ tumors, but fully account for the lower *ITGB4* expression observed in IDH^mut^ gliomas. In fact, *ITGB4* is one of the most downregulated genes in IDH^mut^ 1p/19q co-deleted gliomas compared to IDH^mut^ 1p/19q intact gliomas (*P* = 3.2×10^-39^). In patient-derived glioma cells, we found that integrin β4 is enriched in the stem-like population, and that knockdown of integrin β4 slows glioma growth *in vitro* (*P* < 0.001) and *in vivo (P* = 0.01). Lower *ITGB4* mRNA levels were also associated with longer overall survival in multiple patient cohorts (*P* = 0.0005; *P* < 0.0001). These data suggest that integrin β4 increases glioma malignancy, and that its relative paucity may contribute to the better prognosis in *IDH1/2* mutant, 1p/19q co-deleted oligodendrogliomas.

## Introduction

Gliomas account for 30% of all primary tumors of the central nervous system (CNS) and represent about 80% of malignant CNS tumors (13). Glioblastoma multiforme (GBM) accounts for over 50% of gliomas and carries an extremely poor prognosis, with a median survival of around 14 months even when treating with temozolomide and radiation therapy (57). A key reason for this dismal prognosis is the diffusely infiltrative nature of these tumors, rendering complete surgical excision impossible. Diffuse low and intermediate-grade gliomas have variable outcomes that depend on the underlying genomic alterations present in the tumor, with median survival ranging from 1 to over 12 years (8).

Mutations in *isocitrate dehydrogenases types 1* and *2 (IDH1/2)* are present in the majority of World Health Organization (WHO) grade II and III gliomas, and are also characteristic of secondary GBMs (21, 26, 46, 61). Both enzymes normally oxidize isocitrate to α-ketoglutarate, but point mutations at R132 in *IDH1* or R172 in *IDH2* produce neoenzymatic activity, reducing α-ketoglutarate to D-2-hydroxyglutarate (D2HG) (12, 60). D2HG may contribute to gliomagenesis by promoting global DNA and histone hypermethylation and suppressing cellular differentiation (37). Yet these mutations are powerful favorable prognostic markers, as tumors with mutant *IDH1* or *IDH2* (henceforth referred to as “IDH^mut^”) tend to be much less aggressive compared to their wildtype counterparts (24). However, IDH^mut^ gliomas that also contain 1p/19q co-deletion have an even better prognosis compared to IDH^mut^ gliomas lacking 1p/19q co-deletion: in an analysis of 654 grade II-IV gliomas from The Cancer Genome Atlas (TCGA), the median survival of IDH^mut^, 1p/19q co-deleted gliomas was 134.3 months, compared to 87.5 months for IDH^mut^, 1p/19q intact gliomas and 14.9 months for IDH1^wt^ gliomas (**Supplemental Figure 1**; *P* < 0.0001). The reasons for this survival difference between the major subsets of IDH^mut^ gliomas is incompletely understood.

Integrins are extracellular matrix receptors that regulate a number of diverse cellular processes including survival, motility, angiogenesis and proliferation (19, 41, 58). In addition to their normal functions, integrins can also promote tumor initiation and progression, in part through their ability to enhance receptor tyrosine kinase activity and stimulate proliferative signaling (19). These effects are particularly notable for the integrin α6β4, as it promotes invasion by altering immediate downstream signaling to phosphatidylinositol 3-kinase (PI3K), AKT, MAPK and the Rho small GTPases (19, 36, 42, 43, 53). The integrin α6β4 is comprised of an α and a β subunit that, combined, give the integrin specificity for various laminins. The integrin β4 subunit (referred to herein as integrin β4) has a long cytoplasmic signaling domain that contributes to its ability to promote invasion and metastasis (55). The integrin β4 has also been shown to regulate the expression of genes involved in cell motility, invasion and the metastatic cascade (9). Overexpression of the integrin β4 has been observed in a variety of neoplasms where it is associated with aggressive biology, invasiveness, and poor prognosis (6, 27, 38, 55, 56). Furthermore, integrins facilitate stem cell properties in a number of human malignancies (52), including integrin α6, also known as CD49f, which is the binding partner for integrin β4.

The contribution of integrin β4 to carcinoma progression has been the subject of extensive research in other cancer types; however, expression and function of the integrin β4 has not been studied thoroughly in gliomas. It is not known whether integrin β4 expression is regulated by driver mutations such as in *IDH1/2*, or whether expression of the integrin β4 is influenced by recurrent cytogenetic abnormalities such as 1p/19q co-deletion. As the integrin β4 is known to promote proliferation in other neoplasms, we sought to determine whether integrin β4 contributes to tumor growth and self-renewal in gliomas, and whether differential regulation of *ITGB4* could be responsible for the improved prognosis observed in the IDH^mut^, 1p/19q co-deleted subset. We also sought to determine whether integrin β4 is associated with clinicopathologic variables such as tumor grade and patient survival. Our findings may help explain why IDH^mut^, 1p/19q co-deleted gliomas are even less aggressive than IDH^mut^ gliomas with intact 1p/19q.

## Materials and Methods

### Tissue Microarray Construction

Formalin-fixed, paraffin-embedded (FFPE) glioma tissues were retrieved from the pathology archives at the University of Kentucky. Deidentified tissue microarrays (TMAs) were constructed from the gliomas as previously described (17). A total of 104 cases comprised the TMAs, including 9 nonneoplastic controls (cortical dysplasia), 9 grade II astrocytomas, 11 grade III astrocytomas, 12 anaplastic oligodendrogliomas, 16 grade II oligodendrogliomas, and 47 grade IV GBMs. Treatment and outcomes were obtained via the Kentucky Cancer Registry. Institutional Review Board approval was obtained prior to collecting the archival tissues for TMA construction and patient survival data.

### Immunohistochemistry

Glioma TMA sections (4 μm) were stained using a rat monoclonal primary antibody to the integrin β4 subunit (CD104; clone 439-9B; BD Pharmingen, San Jose, CA) at a concentration of 5μg/ml according to a previously described protocol (11). Integrin β4 expression was scored using a semiquantitative scale as follows: negative (0), minimal staining (1), weakly positive (2), moderately positive (3), strongly and diffusely positive (4). Endothelial cells were used as internal positive controls while non-neoplastic tissue cores served as negative controls. Scoring was done while blinded to WHO grade, IDH^mut^ status, and other clinicopathological variables. Results from each of three tissue cores were averaged together to produce a final score for each tumor.

R132H IDH1 immunohistochemistry of TMAs was performed as previously described (17). Of note, 5 TMA cases were immunonegative for R132H IDH1 but were clinically suspected of having a less common IDH1 or IDH2 mutation based on mutation-associated variables like WHO grade and younger patient age. Of those 5, a single anaplastic astrocytoma turned out to have R132S IDH1 via pyrosequencing, as per standard clinical workup of gliomas.

### Cell culture

The LN18 (CRL-2610) glioma cell line was purchased directly from ATCC and was cultured in RPMI 1640, 10% fetal bovine serum, and 1% penicillin streptomycin, purchased from Life Technologies (Carlsbad, CA). Cells were cultured in humidified conditions at 37°C and 5% CO_2_.

### Cancer stem cell (CSC) derivation and xenograft maintenance

Established patient-derived GBM xenografts were previously reported (3, 20, 51) and were obtained via a material transfer agreement from Duke University (T4121) or the Mayo Clinic (GBM10). Xenografts were originally established under IRB-approved protocols in a deidentified manner using excess tissue taken from consented patients. For experimental studies, GBM cells were dissociated from established xenografts under Cleveland Clinic-approved Institutional Animal Care and Use Committee protocols as described (1). Xenografts were passaged in immune-deficient NOD.Cg-*Prkdc^scid^ II2rg^tm1Wjl^*/SzJ (NSG) mice (obtained from The Jackson Laboratory, Bar Harbor, ME, USA). Freshly dissociated human GBM cells were unilaterally injected into the flank of 6-week-old female mice, and animals were sacrificed by CO_2_ asphyxiation and secondary cervical dislocation when the tumor volume exceeded 5% of the animal’s body weight.

### Tumor dissociation and GBM cell culture

Xenografted tumors were dissected and mechanically dissociated using a papain dissociation kit (Worthington Biochemical Corporation, Lakewood, NJ, USA) as previously described (1, 2, 20, 23, 31, 44). After dissociation, cells were cultured overnight in Neurobasal medium (Life Technologies, Carlsbad, CA, USA) supplemented with B27 (Life Technologies), 1% penicillin/streptomycin (Life Technologies), 1 mM sodium pyruvate, 2 mM L-glutamine, 20 ng/mL EGF (R&D Systems, Minneapolis, MN, USA,), and 20 ng/mL FGF-2 (R&D Systems) in a humidified incubator with 5% CO_2_.

### Self-renewal of integrin β4-high and –low populations using flow cytometry

After dissociation and overnight culture of cells isolated from xenografts, fluorescence-activated cell sorting (FACS) was used to isolate cells based on surface levels of integrin β4 as described for other cell surface receptors (1, 20, 31). Approximately 1×10^6^ cells were filtered through a 70-μm cell strainer, centrifuged, and resuspended in 200 μl of FACS buffer (5% BSA in unsupplemented Neurobasal medium). Two microliters of mouse anti-integrin β4 (Abcam clone M126, ab29042) conjugated to PE/Cy7 (Abcam PE/Cy7^®^ Conjugation Kit, ab102903) was added and incubated for 1 hour in the dark. Cells were then washed three times in unsupplemented Neurobasal medium and resuspended in 500 μl Neurobasal complete. Using DAPI as a control for live cells and an isotype control for signal specificity, the 20% of cells with the highest and 20% of cells with the lowest expression of integrin β4 were plated using a BD FACSAria II at increasing cell concentrations (1, 5, 10, and 20 cells/well) in 96-well plates. After 14 days, the number of wells containing spheres larger than 10 cells across was counted, and the stem cell frequency was calculated using the Walter and Eliza Hall Institute Bioinformatics Division ELDA analyzer (http://bioinf.wehi.edu.au/software/elda/)(25).

### Immunoblotting

Cells were harvested and lysed in RIPA buffer (150 mM NaCl, 0.5 mM EGTA, 0.5% sodium deoxycholate, 0.1% SDS, 1% Triton X-100, 50 mM Tris-HCl pH 7.4, 15 μg/ml protease inhibitor cocktail, 1 mM PMSF, 50 mM NaF and 10 mM sodium pyrophosphate). Total cell lysates (80 μg) were subjected to 8% SDS-PAGE, transferred and immunoblotted with mouse anti-CD104 (integrin β4, BD Bioscience) antibody; β-actin antibodies (Sigma-Aldrich) were used as the loading controls. Data are representative of at least three experiments.

### Quantitative real time PCR (Q-PCR)

Total RNA was extracted using TRizol reagent (Life Technologies/ThermoFisher, Grand Island, NY). Total RNA (1μg) was used to reverse-transcribe into cDNA using the High Capacity cDNA Reverse Transcription Kit (Applied Biosystems/ThermoFisher, Grand Island, NY). Target sequence for ITGB4 shRNA: #4: 5’CGGCTCCTCAGCTACTCCATCCTTC-TCGAGAAGGATGGAGTAGCTGAGGAGTTTTTG and #5: 5’CCGGCCAGCGACTACAC-TATTGGATCTCGAGATCCAATAGTGTAGTCGCT-GGTTTTTG. Expression of *ITGB4* was assessed by Comparative Ct (ΔΔCt) using commercially available probes and Master Mix reagent and performed on a StepOnePlusTM 96-well instrument as described by the manufacturer (Applied Biosystems). The expression level of each gene was normalized by β-actin RNA and reported as a relative level.

### ITGB4 knockdown by shRNA and Xenograft experiments

Patient derived primary GBM stem cells were obtained and isolated as described above. Cells were labeled with CD133 antibody conjugated with APC and sorted to enrich for CD133+ stem cells using FACS. The CD133+ cells were then cultured in Neurobasal Media supplemented with B27 without vitamin A (Invitrogen), 10 ng/mL bFGF (Life Technologies, PHG0026), and 10 ng/mL EGF (Life Technologies, PHG0313). For stable knockdown of *ITGB4* in CD133+ primary GBM stem cells, lentivirus was produced and cells were infected as described previously (54). The cells were then placed under the selection of puromycin. The knockdown efficiency of *ITGB4* was assessed by immunoblotting and Q-PCR. Intracranial tumor implantation was performed by injecting 10 μL (10^5^ cells) into the right cerebral cortex of 5-week-old female NU/J nude mice as previously described (3). Survival was tracked after tumor implantation. Any mouse showing distress, neurological signs (lethargy, hemiparesis, ataxia, seizures) or weight loss of greater than 10% was euthanized in accordance with a protocol approved by the University of Kentucky Institutional Animal Care and Use Committee.

### Bioinformatics analysis of external cohorts

To validate TMA data with external cohorts, the relationship of integrin β4 to tumor type and patient survival was done using the Oncomine microarray database (Oncomine, Compendia Bioscience, Ann Arbor, MI http://www.oncomine.org). Specific correlations between *ITGB4* mRNA and IDH^mut^ status were done via direct query of The Cancer Genome Atlas (TCGA Research Network, http://cancergenome.nih.gov/). Additional survival data in a separate cohort of GBMs was obtained via the National Cancer Institute REpository for Molecular BRAin Neoplasia DaTa (REMBRANDT, https://caintegrator.nci.nih.gov/rembrandt/). TCGA and REMBRANDT datasets were visualized and downloaded using the GlioVis web application (http://gliovis.bioinfo.cnio.es/) (7). Datasets include TCGA grade II-IV gliomas (N = 654) analyzed using RNA-Seq, and REMBRANDT gliomas analyzed using Affymetrix HG U133 v2.0 Plus microarrays.

### Statistical Analysis

Differences between groups were analyzed via two-tailed Student’s t-test, one-way ANOVA with post hoc Tukey’s test, or Kruskal-Wallis test with Dunn’s post hoc test for multiple comparisons where noted. Significance was reached when *P* < 0.05. Survival differences were assessed via log-rank tests for univariate analyses and Cox proportional hazards for multivariate. All univariate statistical analyses were performed using GraphPad Prism, Version 7.01 (Graph Pad Software, La Jolla, CA), and multivariate analyses were done with www.statpages.org.

## Results

To determine the range of integrin β4 expression in gliomas, immunohistochemistry (IHC) for the integrin β4 subunit was performed on a set of glioma TMA sections that included 104 cases (see Materials and Methods). We found that integrin β4 expression was uniformly weak in nonneoplastic control tissues as measured by IHC, but that there was a greater range of staining intensities in grade II-IV gliomas (**Figures 1A-B**). The overall mean staining intensity was significantly higher in grade IV GBMs when compared to grade II-III tumors and nonneoplastic controls (*P* < 0.0001; **Figure 1B**). TCGA analysis confirmed this association, showing elevated *ITGB4* mRNA in GBMs compared to grade II and III tumors, though the difference in means was relatively modest (**Figure 1C**).

**Figure 1:**
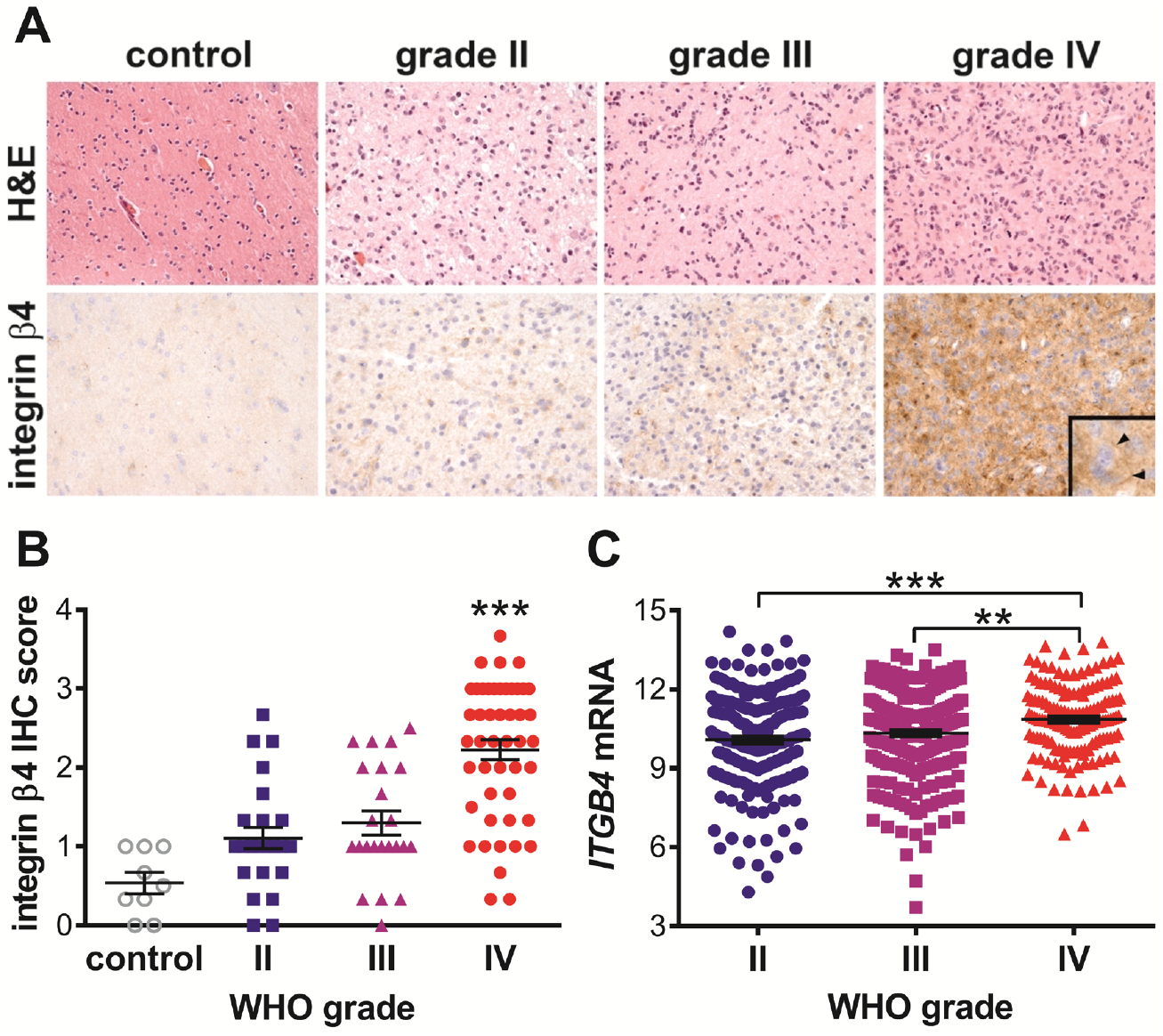
Integrin β4 expression by WHO grade. (A) Representative integrin β4 IHC in WHO grade II-IV gliomas, as well as control nonneoplastic brain tissue. The inset in the grade IV integrin panel highlights membranous staining of integrin β4 in GBM cells (arrowheads). Magnification = 400x for all images. (B) Semiquantitative integrin β4 IHC in control and grade II-IV gliomas. Overall *P* < 0.0001 via one-way ANOVA, *P* < 0.0001 between grade IV GBMs and each other group via post-hoc Tukey’s test. (C) *ITGB4* mRNA in grade II-IV TCGA gliomas. Overall *P* < 0.0001 via one-way ANOVA.

IDH^mut^ has been shown to be a strong favorable prognostic factor in diffusely infiltrative gliomas (24), and high integrin β4 expression can contribute to cancer invasiveness and aggression (55). Therefore, we sought to determine if a correlation between these two molecules existed. Interestingly, a strong inverse relationship between IDH^mut^ and integrin β4 expression was observed in our TMA cohort (*P* < 0.0001; **Figure 2A-B**). Likewise, an analysis of TCGA gliomas showed that IDH^mut^ tumors had lower average *ITGB4* mRNA levels than did IDH^wt^ tumors (*P* = < 0.0001; **Figure 2C**).

**Figure 2:**
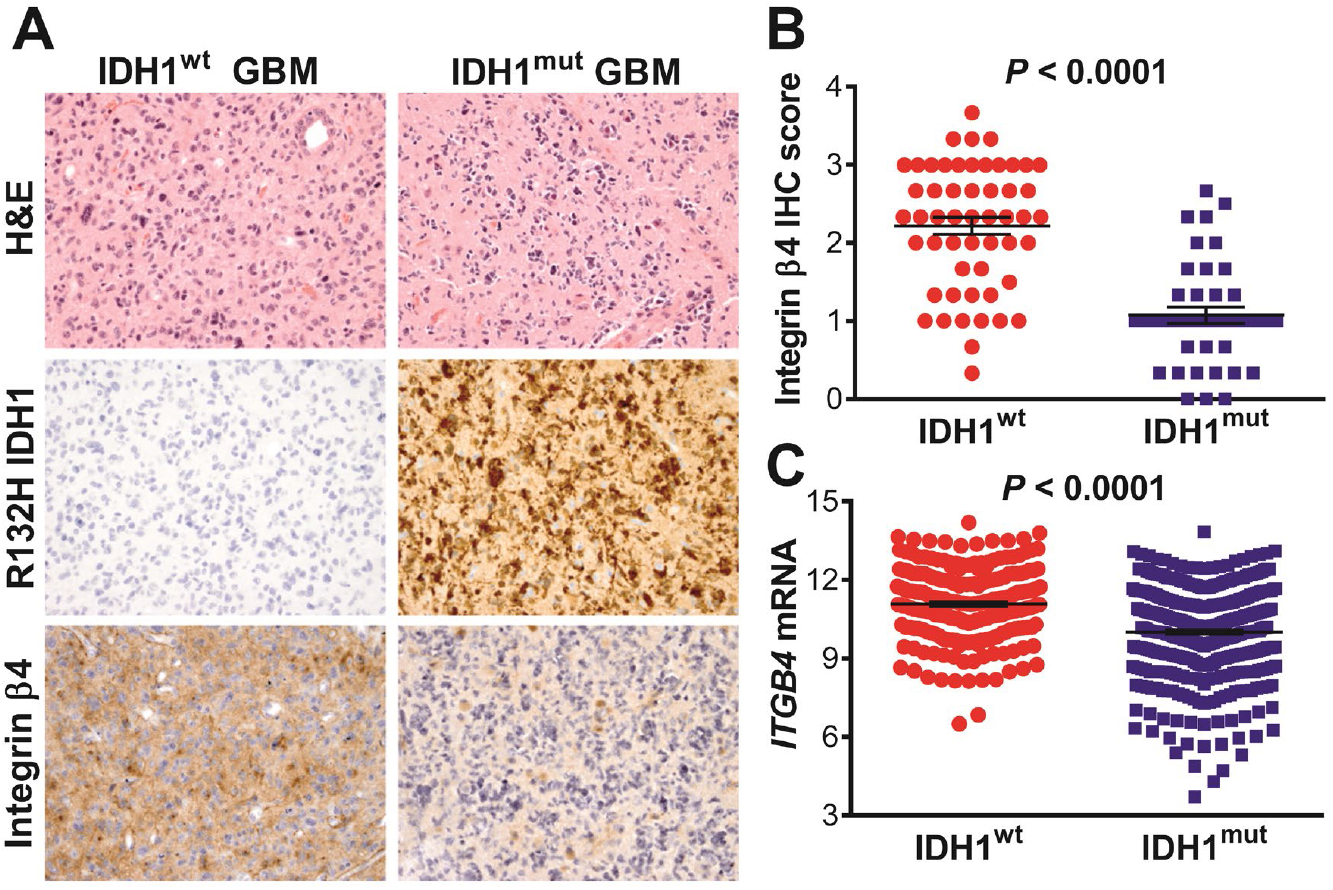
Integrin β4 expression by *IDH* mutation status. (A) Representative photomicrographs of H&Es and IHC for R132H IDH1 and integrin β4 in IDH^wt^ GBM (left panels) and IDH^mut^ GBM (right panels). Magnification = 400x for all images. (B) Semi-quantitative integrin β4 IHC in IDH^wt^ versus IDH^mut^ gliomas. *P* < 0.0001 via two-tailed t-test. (C) *ITGB4* mRNA in IDH^wt^ and IDH^mut^ TCGA gliomas. *P* < 0.0001 via two-tailed t-test.

IDH^mut^ causes global hypermethylation and gene suppression (24). Therefore, we examined CpG sites in the integrin β4 gene *(ITGB4)* and compared its methylation status in IDH^wt^ versus IDH^mut^ tumors. We identified 6 CpG sites in *ITGB4* that are significantly hypermethylated in IDH^mut^ tumors when compared to IDH^wt^ tumors (**Figure 3A, Supplemental Table 1**). Methylation at each of the 6 sites individually correlates with reduced *ITGB4* mRNA (**Figure 3B, Supplemental Table 1**), and 4 of the sites are in the *ITGB4* promoter.

**Figure 3:**
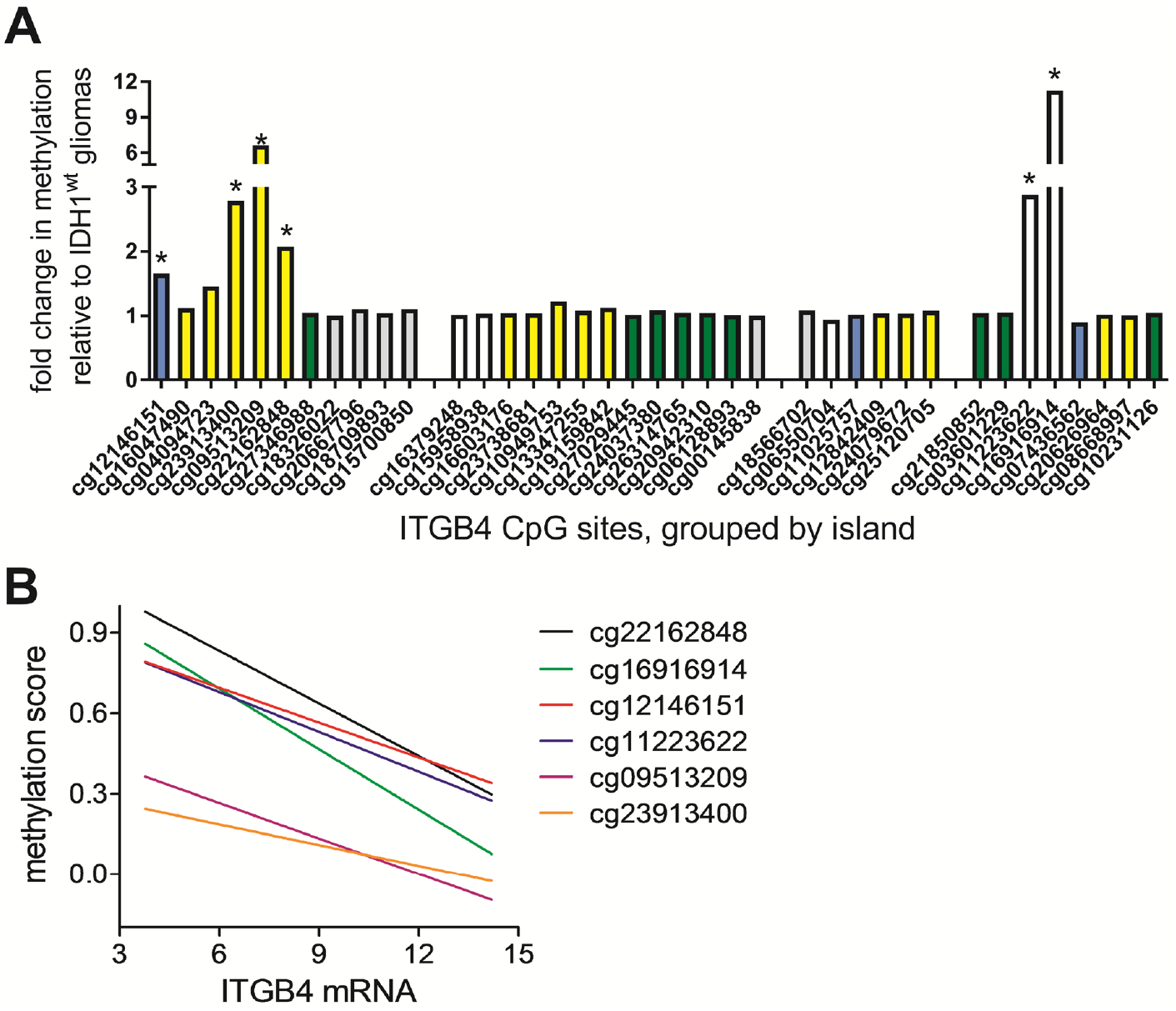
Hypermethylation of *ITGB4* in IDH^mut^ gliomas. (A) Fold difference in methylation of each CpG site in the *ITGB4* gene among IDH^mut^ TCGA gliomas relative to IDH^wt^ TCGA gliomas. (B) Methylation score of significantly hypermethylated CpG sites in the *ITGB4* gene from (A), plotted according to *ITGB4* mRNA levels among all TCGA gliomas.

Gliomas that harbor IDH^mut^ along with 1p/19q co-deletion are histologically oligodendrogliomas and have a particularly good prognosis, even when compared to IDH^mut^ astrocytomas lacking this co-deletion (8, 14). As *ITGB4* promotes tumor invasiveness and aggression, we determined whether differential regulation of *ITGB4* could contribute to the substantially better prognosis observed in this IDH^mut^ subset. By comparing methylation of CpG islands in the *ITGB4* gene between IDH^mut^, 1p/19q intact and IDH^mut^, 1p/19q co-deleted tumors, we found that, while methylation was increased in IDH^mut^, 1p/19q intact astrocytomas relative to IDH^wt^ gliomas, it is even higher (up to 3.7 fold) in the IDH^mut^, 1p/19q co-deleted subset of tumors (**Figure 4A**). Remarkably, only the IDH^mut^, 1p/19q co-deleted oligodendrogliomas showed reduced *ITGB4* mRNA relative to IDH^wt^ gliomas, not the IDH^mut^, 1p/19q intact astrocytomas (*P* < 0. 0001; **Figure 4B**). Among 20,530 genes analyzed in the TCGA via RNA-Seq, *ITGB4* is the 56^th^ most downregulated gene in IDH^mut^ 1p/19q co-deleted gliomas relative to IDH^mut^ 1p/19q intact gliomas *P*=3.2×10^-39^, **Supplemental Table 2**). We also compared integrin β4 IHC between the IDH^mut^ and IDH^mut^, 1p/19q co-deleted subsets in the TMA. However, we did not detect a significant difference in expression in this cohort (*P* = 0.60; **Figure 4C**); the TMA cohort was likely underpowered to detect such a difference.

**Figure 4:**
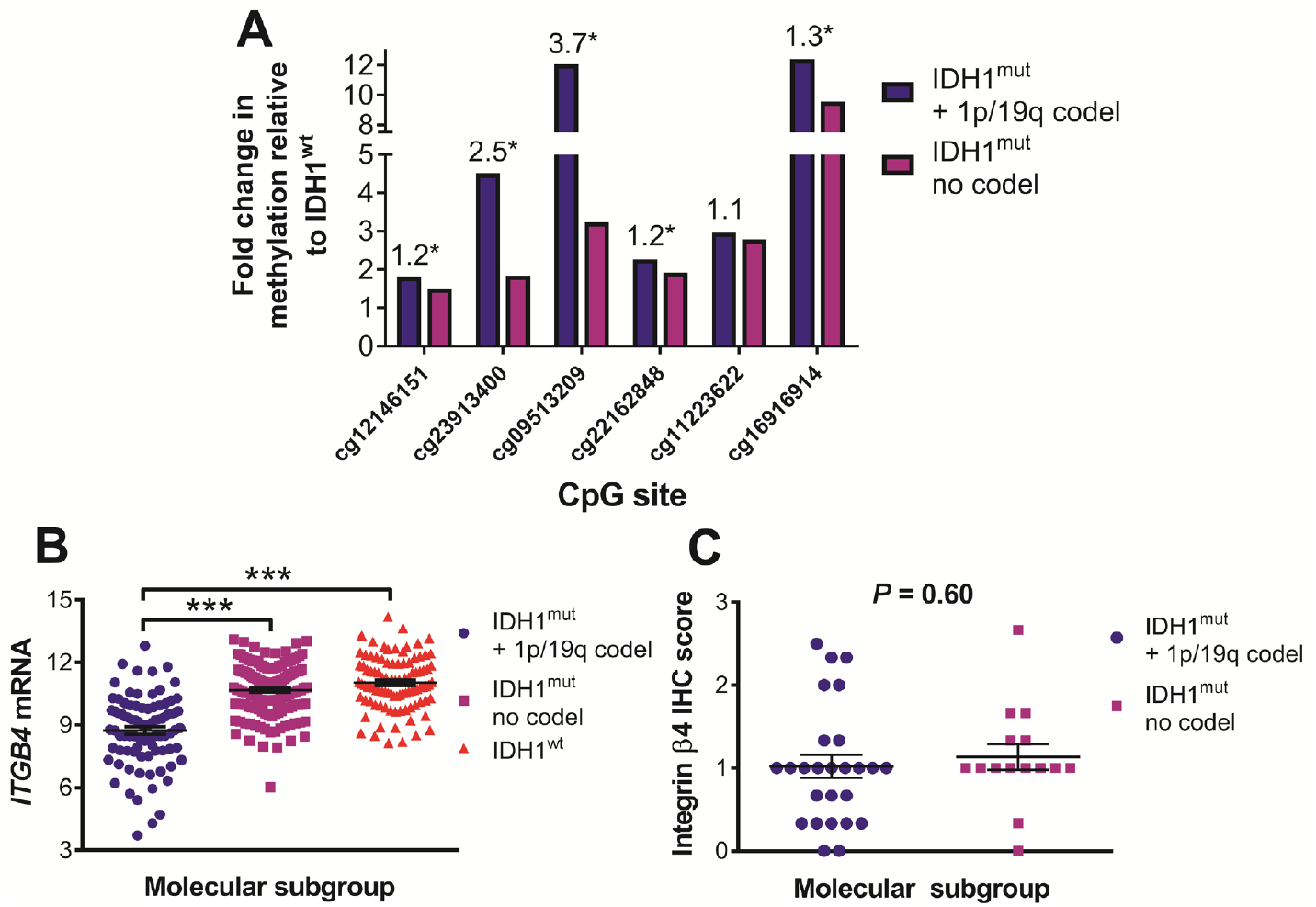
Differential *ITGB4* expression and methylation between major subtypes of IDH^mut^ gliomas. (A) Methylation of CpG sites in *ITGB4* gene in IDH^mut^, 1p/19q co-deleted oligodendrogliomas compared with 1p/19q intact IDH^mut^ astrocytomas. (B) *ITGB4* mRNA levels IDH^mut^, 1p/19q co-deleted oligodendrogliomas compared with 1p/19q intact IDH^mut^ astrocytomas (*P* < 0.0001) and IDH^wt^, 1p/19q intact GBM (*P* < 0.0001). (C) Integrin β4 IHC semi-quantitative score between IDH^mut^, 1p/19q co-deleted gliomas and those with intact 1p/19q (*P* = 0.60 via two-tailed t test) in our institutional TMA cohort.

Glioma stem cells (GSCs) represent a subset of cells that contribute to glioma growth and self-renewal, and are believed to promote therapy resistance and evasion of anticancer host immunity (32). Previous studies have demonstrated that integrins can mediate cancer stem-cell properties (52), and this led us to investigate the relationship between integrin β4 expression and self-renewal in patient-derived, CD133+ IDH^wt^ primary GBM cells. Tumors were sorted for integrin β4 surface expression using FACS, and limiting dilution analysis was used to estimate the stem cell frequency of cells isolated from freshly dissociated GBM xenografts. We found that across multiple specimens, integrin β4-high cells had higher stem cell frequencies than was observed in integrin β4-low cells (**Figure 5A-D**). To further investigate the role of integrin β4 in glioma growth, we performed loss-of-function studies in cell lines and in an intracranial patient-derived xenograft model of glioblastoma. Efficient knockdown of *ITGB4* (**Figure 6A-B**) in LN18 cells dramatically decreased *in vitro* glioma growth (*P* < 0.001; **Figure 6C**), and significantly extended the survival of mice engrafted with patient-derived, CD133+ IDH^wt^ primary GBM cells (median survival 24.5 days vs. 33 days; *P* = 0.01; **Figure 6D-E**).

**Figure 5:**
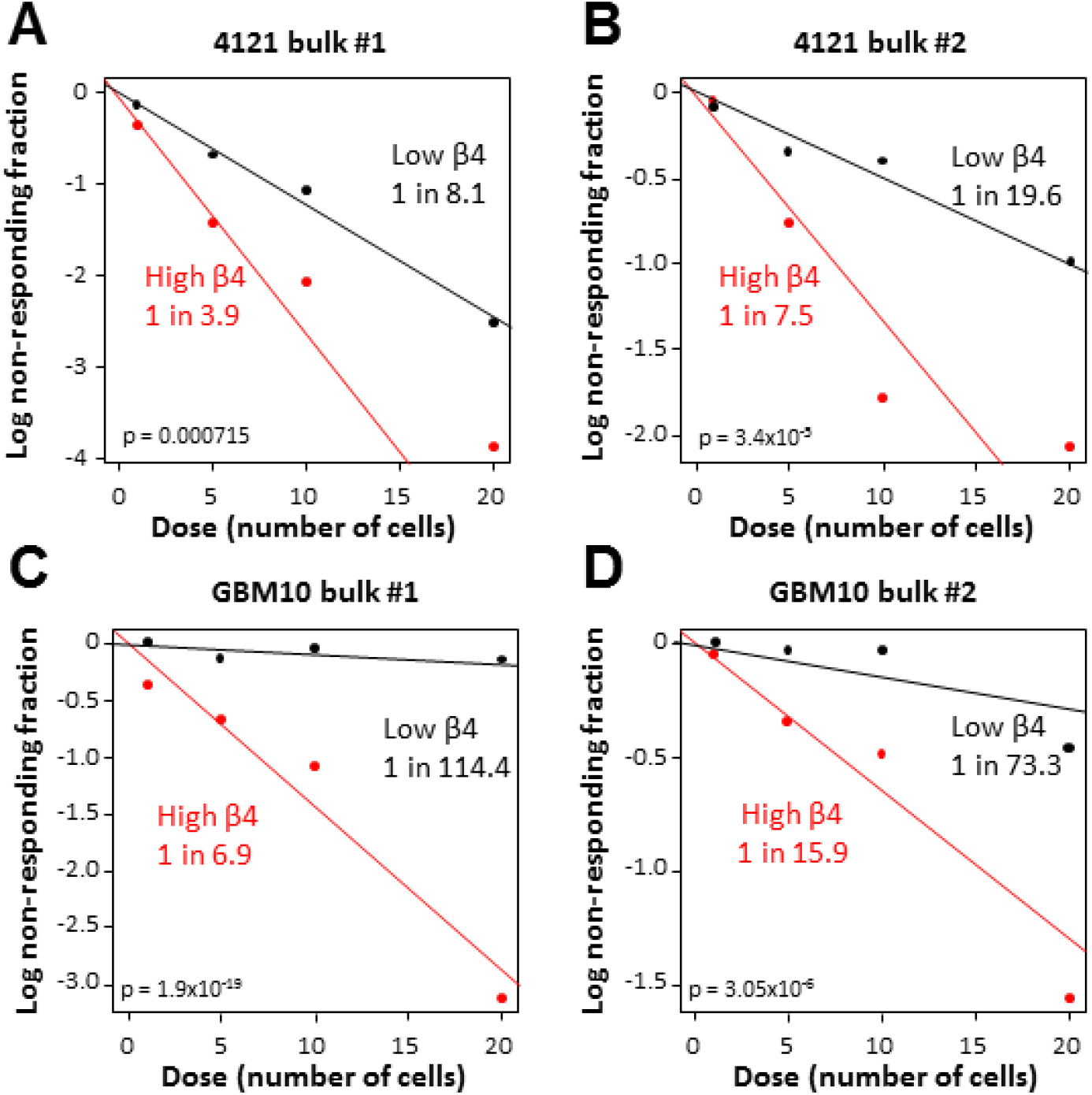
Integrin β4 in glioma stem cells. (A-D) Limiting dilution analysis was utilized to estimate the stem cell frequency of cells isolated from freshly dissociated GBM xenografts. The 20% of cells with the highest and lowest expression of integrin β4 were plated in a limiting dilution manner, and the number of wells containing spheres were counted after 14 days to generate stem cell frequencies.

**Figure 6:**
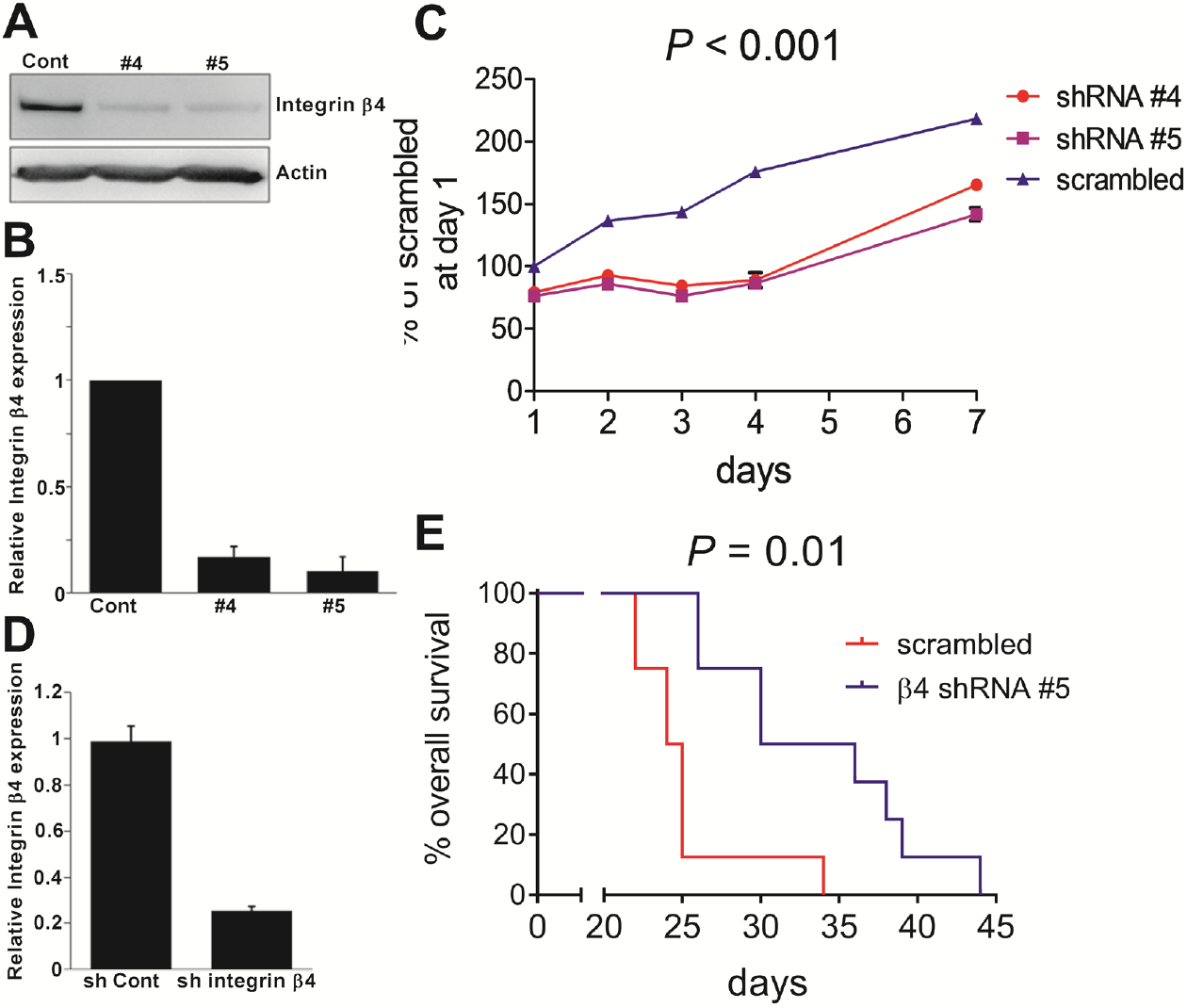
*ITGB4* knockdown suppresses glioma growth. (A-C) Effect of *ITGB4* knockdown on *in vitro* growth of LN-18 cells. (D-E) Effect of *ITGB4* knockdown on *in vivo* growth of patient-derived, CD133+ primary GBM cells xenografted into the brains of immunocompromised mice.

To date, no data on integrin β4 and outcome in gliomas has been published. In our TMA cohort, elevated integrin β4 expression was significantly associated with shorter overall survival among all grade II-IV gliomas (median survival not reached for integrin β4 IHC score <1; 63.5 months for 1-1.9; 18.1 months for 2+, *P* = 0.0005; **Figure 7A**). We also examined the relationship between *ITGB4* mRNA expression and patient survival in the TCGA glioma cohort (*N* = 656), where we found that increasing levels of *ITGB4* mRNA correlated with worse overall survival (**Figures 7B**). Moreover, in the TCGA glioma cohort, *ITGB4* expression quartile was found to be an independent prognostic factor, even when including variables such as IDH^mut^ and 1p/19q co-deletion status (**Table 1**).

**Table 1:**
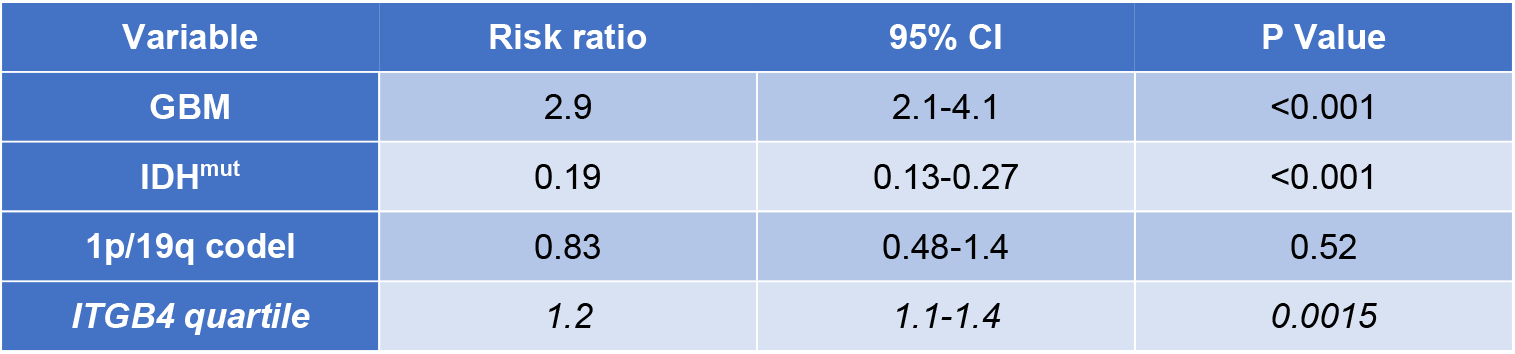
Multivariable analysis of TCGA gliomas according to specific prognostic markers. Cox proportional hazards modeling of TCGA gliomas, including GBM, IDH^mut^ status, 1p/19q codeletion, and *ITGB4* mRNA quartile (see **Figure 7B**) as variables.

**Figure 7:**
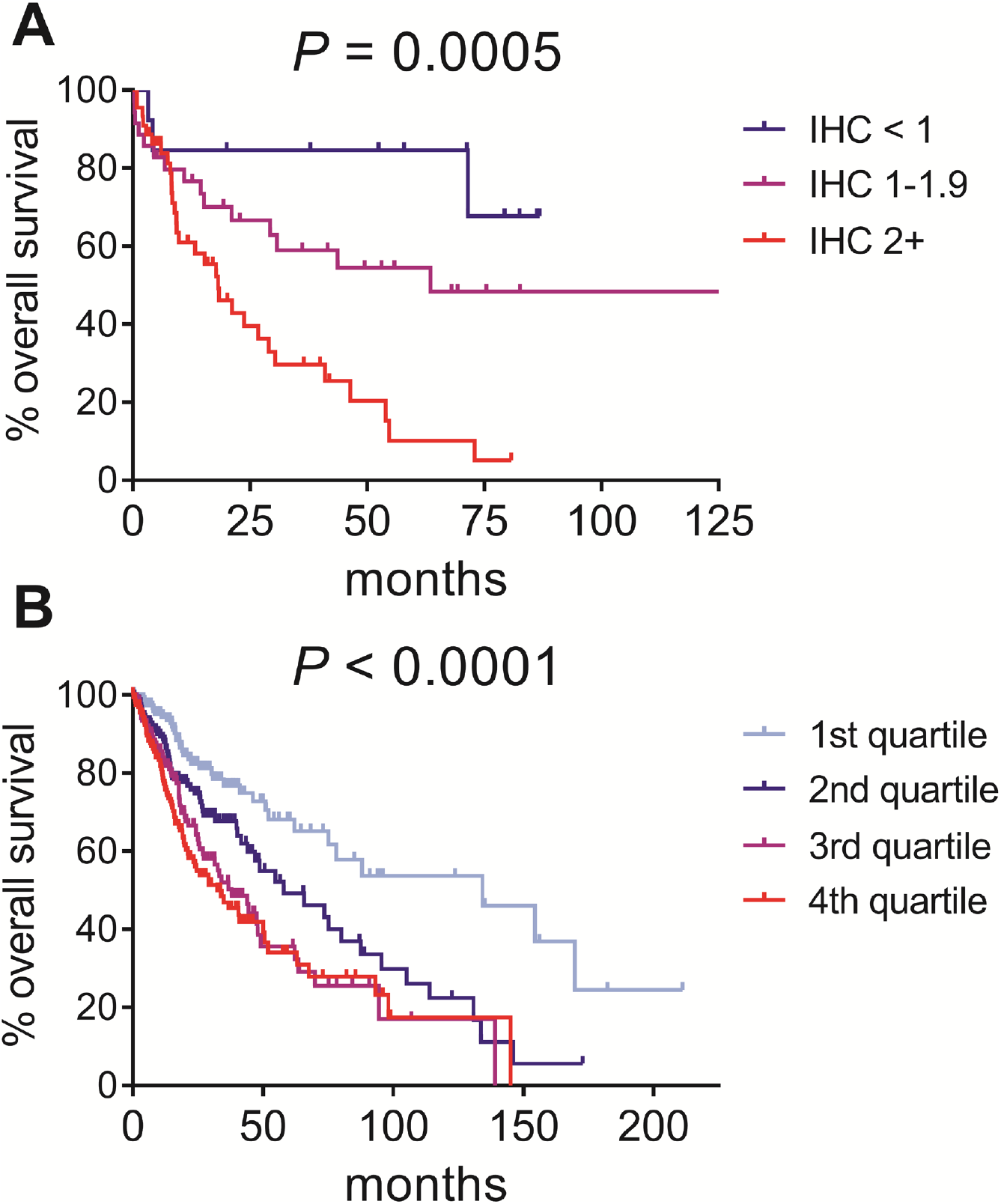
Integrin β4 and survival in gliomas. (A) Overall survival according to integrin β4 IHC in all grade II-IV gliomas from our institutional TMA cohort (*P* = 0.0005 via log-rank test). (B) Overall survival according to *ITGB4* mRNA in grade II-IV TCGA gliomas *(N* = 656, *P* < 0.0001 via log-rank test).

## Discussion

Our results demonstrate that integrin β4 is highly expressed in GBMs compared to lower grade gliomas and non-neoplastic brain tissue. We found that elevated integrin β4 expression is an adverse prognostic marker, which is consistent with work done in other cancers showing that integrin β4 overexpression imparts more aggressive behavior (55). We provide evidence that integrin β4 is enriched in the stem-like subset of glioma cells and that it contributes to glioma growth and self-renewal. We find that integrin β4 expression is inversely correlated with IDH^mut^ status, and that methylation and expression of *ITGB4* differ according to 1p/19q co-deletion status. Given the role of integrin β4 in tumor malignancy, these findings to help explain why IDH^mut^, 1p/19q co-deleted oligodendrogliomas generally are even less aggressive than IDH^mut^ astrocytomas with intact 1p/19q (50, 62).

Global hypermethylation contributes to tumorigenesis by virtue of its ability to suppress cellular differentiation, yet the effects of generalized hypermethylation can eventually lead to changes that inhibit tumor malignancy (24). For example, IDH^mut^ is associated with hypermethylation of the *MGMT* promoter and suppression of MGMT (40), thereby reducing the ability of the tumor to repair DNA damage caused by alkylating chemotherapeutic agents such as temozolomide (22). We find that methylation of CpG islands in the *ITGB4* promoter is significantly higher in IDH^mut^ tumors with 1p/19q co-deletion, and that this hypermethylation is associated with a concomitant reduction in *ITGB4* gene expression. In agreement with these findings, epigenetic regulation of integrin β4 expression through promoter hypermethylation and histone modification has been previously described (35).

Mutations in *IDH1* are known to promote global DNA and histone methylation, although the exact genes that are methylated can differ between tumor types (18, 28). We find that while *ITGB4* promoter methylation is generally higher in IDH^mut^ gliomas, this phenomenon is mostly due to the 1p/19q co-deleted subset of IDH^mut^ tumors. At this point, it is unclear what mechanism is responsible for this observation. However, in one study of IDH^mut^ gliomas, *MGMT* promoter methylation was even higher in 1p/19q co-deleted oligodendrogliomas compared to 1p/19q intact astrocytomas, suggesting that secondary genetic alterations may affect the type and degree of IDH^mut^-induced methylation (34). Genes such as *FGFR2* and *TNFRFS1B* have been shown to be differentially methylated in 1p/19q co-deleted oligodendrogliomas compared to IDH^mut^ astrocytomas (28). Furthermore, secondary mutations, such as in *ATRX* and *TP53* in 1p/19q intact gliomas, and *CIC* and *FUBP1* in 1p/19q co-deleted gliomas, may also influence which genes are hypermethylated in IDH^mut^ tumors.

Our results provide further evidence for the importance of integrins in the GSC population, as we found that integrin β4 is enriched in CD133+ glioma stem cells and that it contributes to glioma self-renewal. A variety of stem cell populations use integrins to associate with extracellular matrix proteins, a relationship that is important for stem cell homing and niche interaction (10). The integrin β4 interacts with laminins in the extracellular matrix (55), and laminin-integrin signaling plays a role in the development of neural stem cells (33). In the neoplastic setting, high expression of integrin α6 is a characteristic of GBM stem cells, and its interaction with laminin α2 in the extracellular matrix mediates stem cell properties like selfrenewal and tumor formation (29, 30). In glioblastoma, the integrins α3, α6 and β1 cooperate with CD151 to regulate stemness and self-renewal (59). Integrin β4 is also a marker for cancer stem cells in other tumor types, and a recent paper demonstrated that integrin β4 identifies a stem-cell enriched population of partially mesenchymal cells in patients with triple negative breast carcinoma (6).

In addition to their ability to mediate stem-cell properties, integrins have been shown to contribute to other aspects of glioma biology. Integrin β8 is expressed in gliomas, positively correlates with overexpression of Egfr and Pdgfr-α, and may promote Fak and Erk activity in regions rich in extracellular matrix (49). The αvβ3 and αvβ5 integrins have been implicated in tumor-induced angiogenesis (16), which is a common feature of GBMs (15). The αvβ3 and αvβ5 integrins increase with increasing histologic grade in gliomas, and tumor cells at the periphery of high-grade gliomas have higher αvβ3 expression (4). Attempts to target αvβ3 and αvβ5 in GBM led to multiple clinical trials with cilengitide, a small molecule inhibitor of the αvβ3, αvβ5 and α5β1 integrins (39).

To date, few studies have examined integrin β4 expression in gliomas. However, expression of the integrin β4 has been identified in astrocytomas, oligodendrogliomas, and GBMs, as well as in reactive astrocytes (5, 45, 47). One study found that the integrin β4 was expressed in a mouse model of GBM, that integrin β4 was present early in tumor development, and that expression was more intense within the neoplastic cells of proliferative centers (48).

In summary, we demonstrate for the first time that expression of the β4 integrin is specifically decreased in IDH^mut^, 1p/19q co-deleted oligodendrogliomas, and that high integrin β4 correlates with poor prognosis in glioma patients. Further work is needed to clarify the mechanisms by which IDH^mut^ and 1p/19q co-deletion may alter the expression of cellular adhesion molecules, and to examine the role of integrin α6β4 in gliomagenesis and invasion. Gaining a deeper understanding of why IDH^mut^ gliomas, especially those with 1p/19q codeletion, are less aggressive than their IDH’“‘ counterparts will not only help develop ways of targeting IDH^mut^ gliomas, but also of making IDH^wt^ tumors behave more like IDH^mut^ tumors.

## Acknowledgments

This work was supported by National Institutes of Health R01 NS102669 (CH), K08 CA155764 (CH), National Institute of General Medical Sciences RR020171 (CH), The Peter and Carmen Lucia Buck Training Program in Translational Clinical Oncology (CH), and the University of Kentucky College of Medicine Physician Scientist Program (CH); National Institutes of Health T32 CA160003 (RLS); and National Institutes of Health R01 CA109136 (MC and KLO). These studies were supported by a NIH K99/R00 Pathway to Independence Award (CA157948) to JDL. Work in the Lathia lab is also supported the National Institutes of Health (R01 NS083629), the Lerner Research Institute, Voices Against Brain Cancer, the Ohio Cancer Research Associates, V Scholar Award from the V Foundation for Cancer Research, Grant IRG-91-022-18 to the Case Comprehensive Cancer Center from the American Cancer Society, and the Cleveland Clinic Product Development Fund.

The Markey Biospecimen and Tissue Procurement (BSTP) Shared Resource Facility facilitated the construction of tissue microarrays and immunohistochemical studies. Special thanks to Dana Napier for her excellent histologic expertise. The BSTP is supported by the University of Kentucky Markey Cancer Center (P30CA177558). The Northwestern Nervous System Tumor Bank is supported by the P50CA221747 SPORE for Translational Approaches to Brain Cancer.

## Conflict of Interest

The authors report no conflicts of interest.

## SUPPLEMENTAL DATA

**Supplemental Figure 1:**
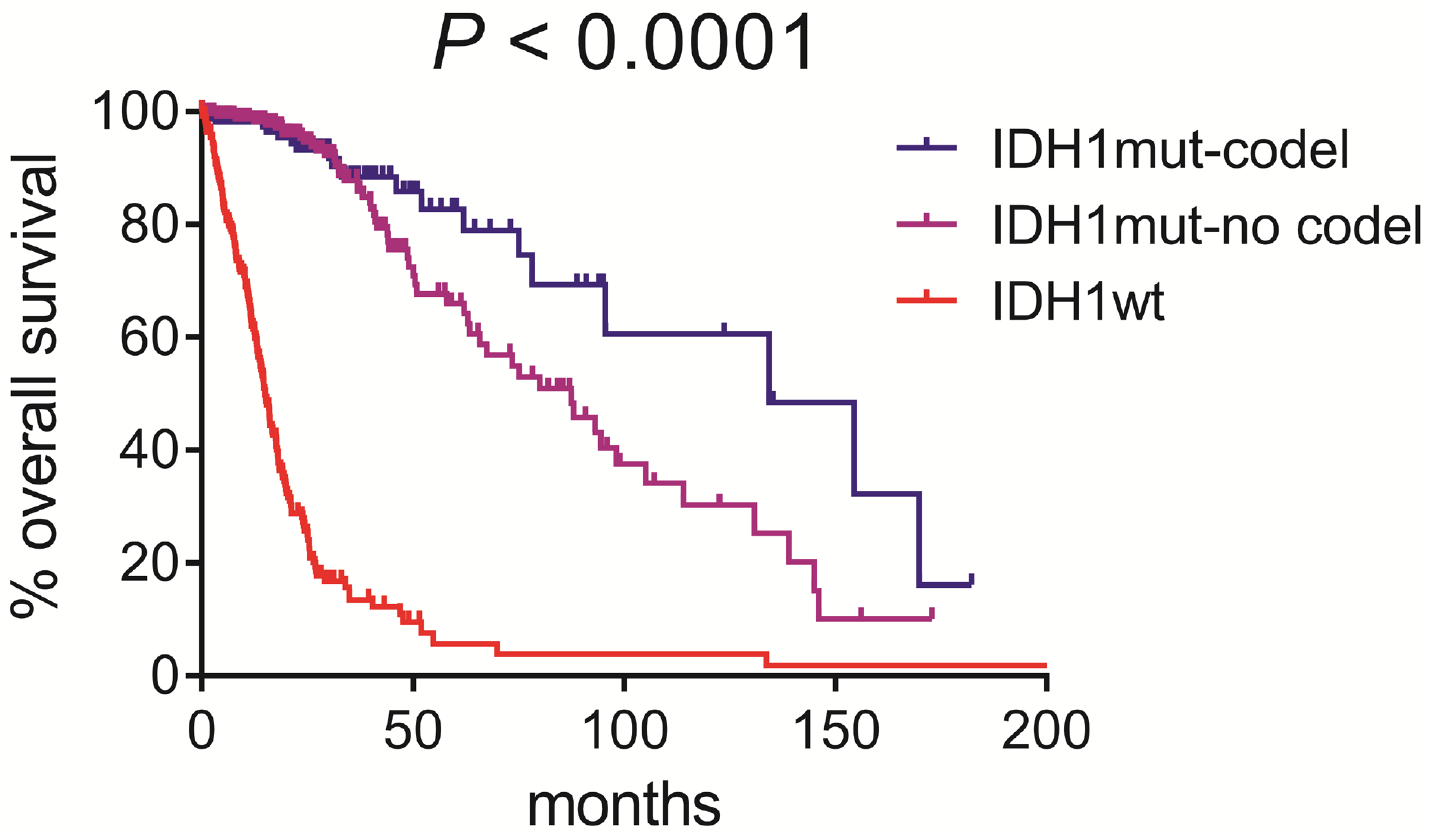
Molecular subtype and overall survival in TCGA gliomas. In a TCGA cohort (*N* = 654) the median survival for patients with IDH^mut^, 1p/19q co-deleted gliomas was significantly higher (134.3 months) when compared to that of patients with IDH1mut, 1p/19q intact gliomas (87.5 months) and IDH^wt^ gliomas (14.9 months; *P* < 0.0001).

**Supplemental Table 1:** Methylation of CpG sites within *ITGB4* gene in TCGA gliomas. See Figure 3 in main text.

| ID# | position relative<br>to island | coordinate | relation to ITGB4<br>coding region | IDH1wt | IDH1mut | fold change | P value | slope | 95% CI | R square |
| --- | --- | --- | --- | --- | --- | --- | --- | --- | --- | --- |
| cg12146151 | N_Shore | 73716945 | TSS1500 | 0.35 | 0.58 | 1.66 | 7.60572E-41 | -0.04328 ± 0.005710 | -0.05447 to -0.03208 | 0.15 |
| cg16047490 | Island | 73717197 | TSS1500 | 0.02 | 0.02 | 1.12 | 0.021987713 |  |  |  |
| cg04094723 | Island | 73717212 | TSS1500 | 0.02 | 0.03 | 1.46 | 0.002947797 |  |  |  |
| cg23913400 | Island | 73717552 | 5'UTR | 0.03 | 0.09 | 2.78 | 3.47511E-15 | -0.02593 ± 0.002361 | -0.03056 to -0.02130 | 0.26 |
| cg09513209 | Island | 73717562 | 5'UTR | 0.02 | 0.10 | 6.64 | 7.4081E-14 | -0.04422 ± 0.003424 | -0.05093 to -0.03751 | 0.33 |
| cg22162848 | Island | 73717736 | 5'UTR | 0.31 | 0.65 | 2.07 | 1.8728E-51 | -0.06526 ± 0.007446 | -0.07986 to -0.05067 | 0.19 |
| cg27346988 | S_Shore | 73719693 | 5'UTR | 0.77 | 0.80 | 1.04 | 7.64758E-09 |  |  |  |
| cg18326022 | S_Shelf | 73719885 | 5'UTR | 0.97 | 0.97 | 1.00 | 0.038764352 |  |  |  |
| cg20667796 | S_Shelf | 73720234 | 5'UTR | 0.70 | 0.78 | 1.10 | 1.16883E-17 |  |  |  |
| cg18709893 | S_Shelf | 73720591 | 5'UTR | 0.83 | 0.86 | 1.04 | 4.85836E-09 |  |  |  |
| cg15700850 | S_Shelf | 73720840 | Body | 0.77 | 0.84 | 1.10 | 3.22756E-25 |  |  |  |
| cg16379248 | N_Shelf | 73723681 | Body | 0.92 | 0.93 | 1.01 | 0.000561755 |  |  |  |
| cg15958938 | Island | 73726239 | Body | 0.90 | 0.92 | 1.03 | 5.12804E-06 |  |  |  |
| cg16603176 | Island | 73726241 | Body | 0.88 | 0.91 | 1.04 | 9.10955E-07 |  |  |  |
| cg23738681 | Island | 73726337 | Body | 0.88 | 0.91 | 1.03 | 7.78461E-17 |  |  |  |
| cg10949753 | Island | 73726372 | Body | 0.77 | 0.95 | 1.22 | 9.79778E-37 |  |  |  |
| cg13347255 | Island | 73726430 | Body | 0.81 | 0.88 | 1.08 | 7.56851E-19 |  |  |  |
| cg19159842 | S_Shore | 73727435 | Body | 0.73 | 0.82 | 1.13 | 4.48068E-26 |  |  |  |
| cg27029445 | S_Shore | 73727713 | Body | 0.93 | 0.94 | 1.01 | 6.73622E-05 |  |  |  |
| cg24037380 | S_Shore | 73727879 | Body | 0.84 | 0.92 | 1.09 | 3.60522E-11 |  |  |  |
| cg26314765 | S_Shore | 73727957 | Body | 0.94 | 0.98 | 1.04 | 2.82777E-07 |  |  |  |
| cg20942310 | S_Shore | 73728061 | Body | 0.85 | 0.89 | 1.04 | 1.18098E-07 |  |  |  |
| cg06128893 | S_Shelf | 73729655 | Body | 0.97 | 0.98 | 1.01 | 0.000958105 |  |  |  |
| cg00145838 | S_Shelf | 73729676 | Body | 0.99 | 0.99 | 1.00 | 0.130023661 |  |  |  |
| cg18566702 | N_Shore | 73736931 | Body | 0.73 | 0.79 | 1.08 | 3.4272E-14 |  |  |  |
| cg06550704 | Island | 73738580 | Body | 0.87 | 0.82 | 0.94 | 0.000728413 |  |  |  |
| cg11025757 | Island | 73738713 | Body | 0.97 | 0.98 | 1.02 | 0.002526145 |  |  |  |
| cg12842409 | Island | 73738809 | Body | 0.91 | 0.94 | 1.03 | 2.54487E-07 |  |  |  |
| cg24079672 | S_Shore | 73739840 | Body | 0.96 | 0.99 | 1.03 | 2.13744E-05 |  |  |  |
| cg25120705 | S_Shore | 73739874 | Body | 0.90 | 0.97 | 1.08 | 1.85209E-21 |  |  |  |
| cg21850852 | N_Shelf | 73746186 | Body | 0.06 | 0.06 | 1.04 | 0.440958354 |  |  |  |
| cg03601229 | N_Shore | 73748344 | Body | 0.85 | 0.89 | 1.05 | 1.8649E-12 |  |  |  |
| cg11223622 | Island | 73749654 | Body | 0.20 | 0.58 | 2.88 | 2.9742E-110 | -0.04934 ± 0.006558 | -0.06219 to -0.03649 | 0.14 |
| cg16916914 | Island | 73749694 | Body | 0.04 | 0.51 | 11.29 | 1.06347E-75 | -0.07510 ± 0.009054 | -0.09285 to -0.05735 | 0.17 |
| cg07436562 | S_Shore | 73750299 | Body | 0.40 | 0.36 | 0.90 | 8.81742E-05 |  |  |  |
| cg20626964 | S_Shelf | 73753034 | Body | 0.80 | 0.82 | 1.02 | 4.15866E-06 |  |  |  |
| cg08668997 | S_Shelf | 73753076 | Body | 0.98 | 0.99 | 1.01 | 0.000232337 |  |  |  |
| cg10231126 | S_Shelf | 73753800 | 3'UTR | 0.85 | 0.89 | 1.04 | 6.75818E-08 |  |  |  |

**Supplemental Table 2:**
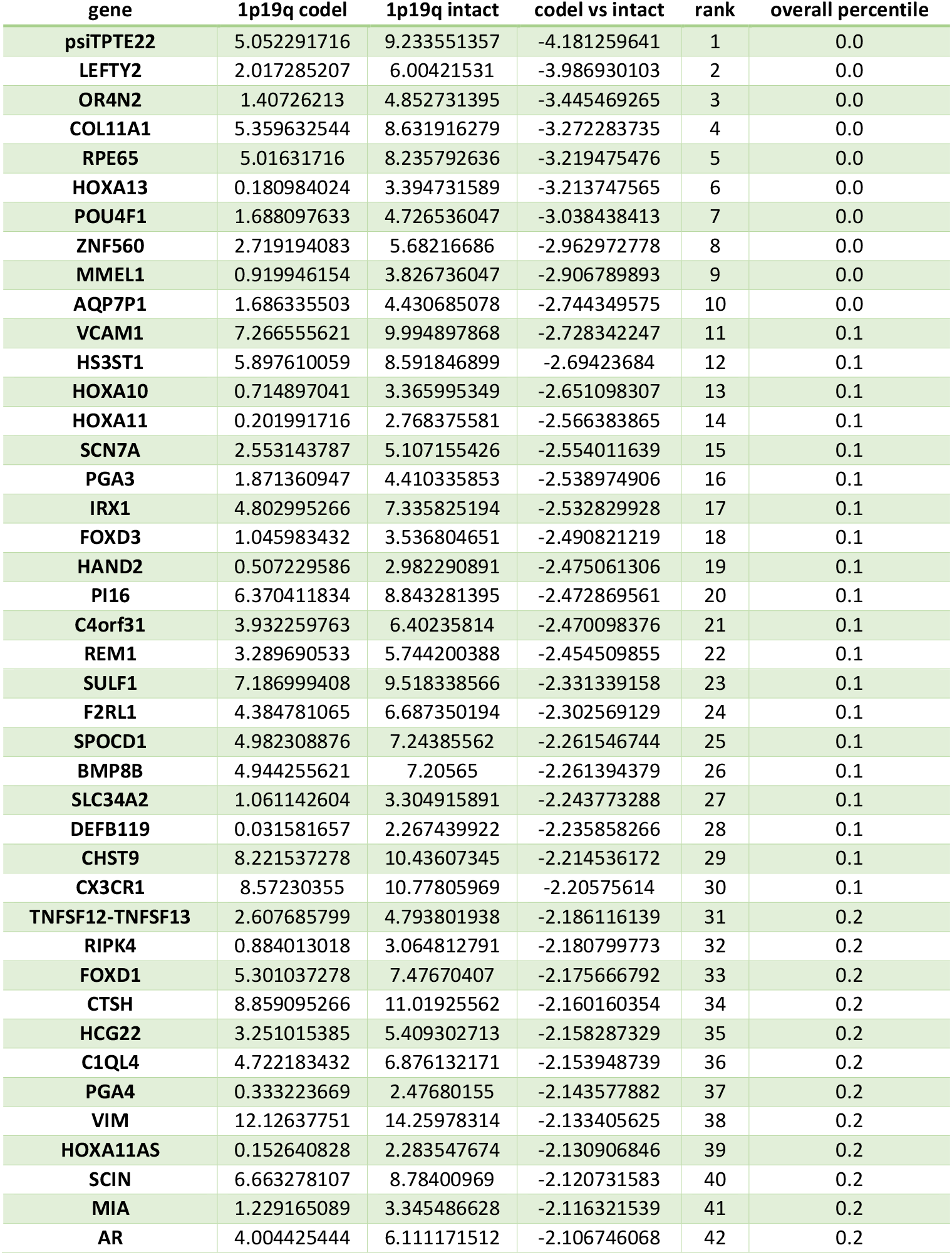
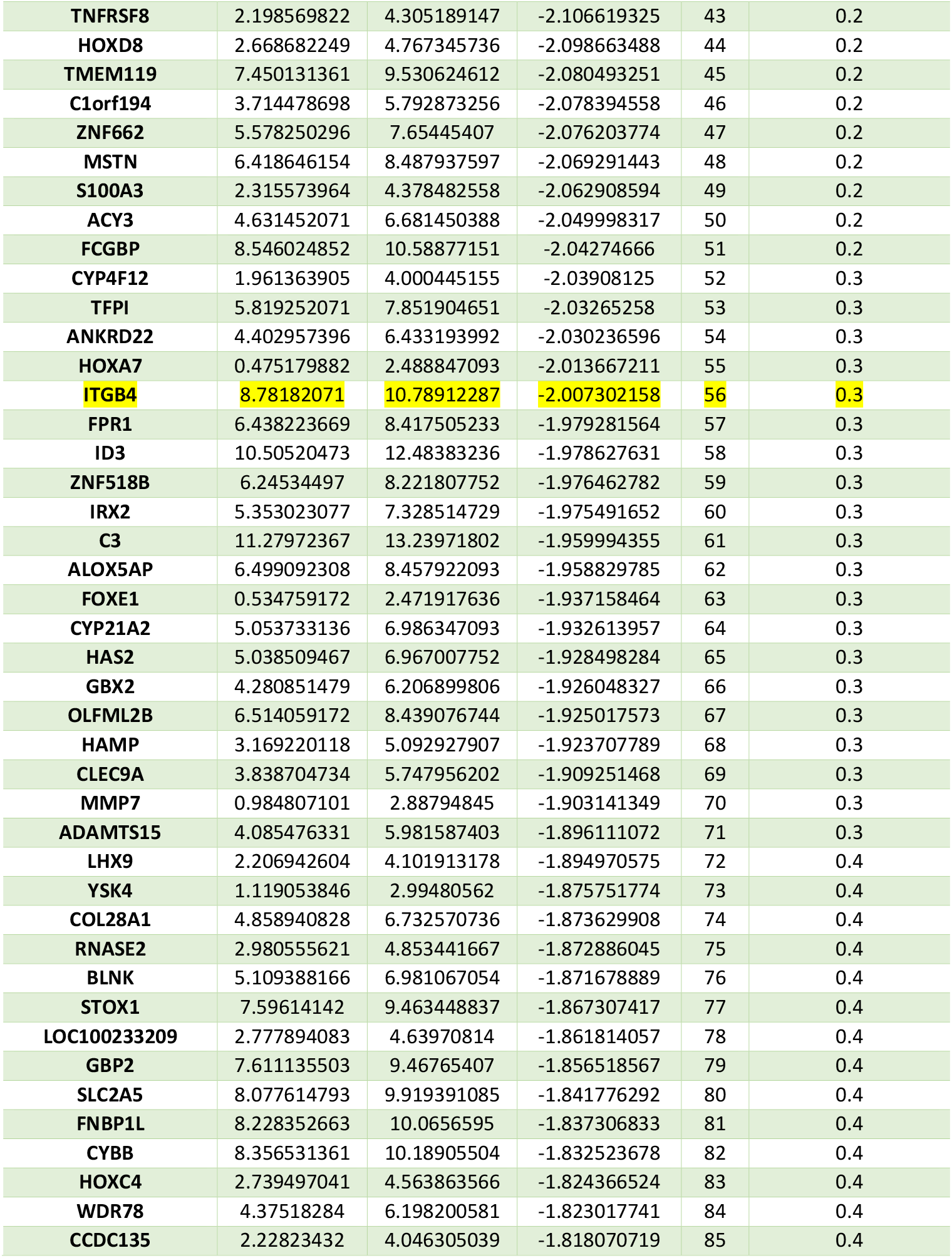
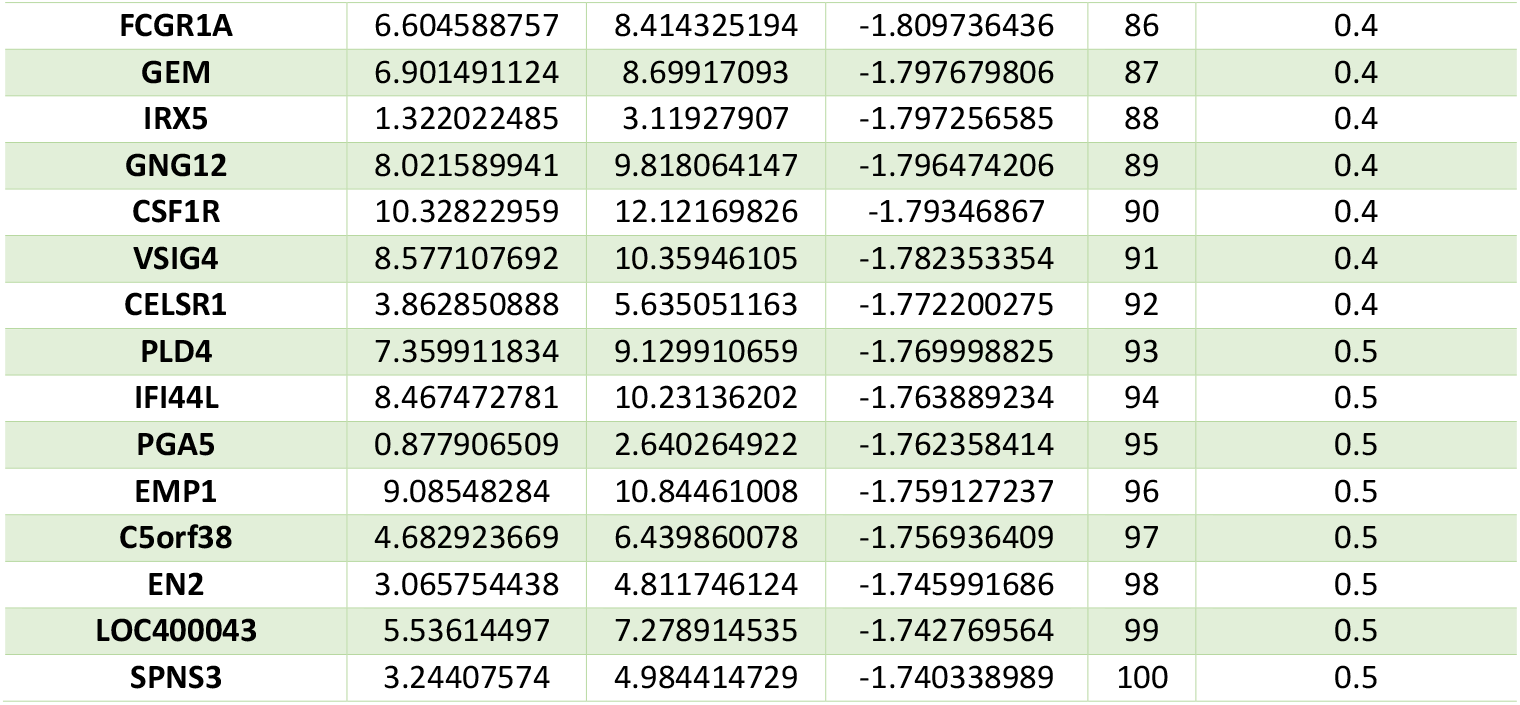
List of the 100 most downregulated genes in IDH^mut^ 1p/19q co-deleted oligodendrogliomas relative to IDH^mut^ 1p/19q intact astrocytomas. Among 20,530 genes analyzed in TCGA gliomas via RNA-Seq, *ITGB4* is the 56^th^ most downregulated gene in IDH^mut^ 1p/19q co-deleted oligodendrogliomas relative to IDH^mut^ 1p/19q intact astrocytomas (*P* = 3.2×10^-39^).

## References

1. Alvarado AG, Thiagarajan PS, Mulkearns-Hubert EE, Silver DJ, Hale JS, Alban TJ, et al. Glioblastoma Cancer Stem Cells Evade Innate Immune Suppression of Self-Renewal through Reduced TLR4 Expression. Cell stem cell. 2016.

2. Alvarado AG, Turaga SM, Sathyan P, Mulkearns-Hubert EE, Otvos B, Silver DJ, et al. Coordination of self-renewal in glioblastoma by integration of adhesion and microRNA signaling. Neuro-oncology. 2016;18(5):656–66.

3. Bao S, Wu Q, McLendon RE, Hao Y, Shi Q, Hjelmeland AB, et al. Glioma stem cells promote radioresistance by preferential activation of the DNA damage response. Nature. 2006;444(7120):756–60.

4. Bello L, Francolini M, Marthyn P, Zhang J, Carroll RS, Nikas DC, et al. Alpha(v)beta3 and alpha(v)beta5 integrin expression in glioma periphery. Neurosurgery. 2001;49(2):380–9; discussion 90.

5. Belot N, Rorive S, Doyen I, Lefranc F, Bruyneel E, Dedecker R, et al. Molecular characterization of cell substratum attachments in human glial tumors relates to prognostic features. Glia. 2001;36(3):375–90.

6. Bierie B, Pierce SE, Kroeger C, Stover DG, Pattabiraman DR, Thiru P, et al. Integrin-beta4 identifies cancer stem cell-enriched populations of partially mesenchymal carcinoma cells. Proceedings of the National Academy of Sciences of the United States of America. 2017;114(12):E2337–E46.

7. Bowman RL, Wang Q, Carro A, Verhaak RG, Squatrito M. GlioVis data portal for visualization and analysis of brain tumor expression datasets. Neuro-oncology. 2017;19(1):139–41.

8. Cancer Genome Atlas Research N, Brat DJ, Verhaak RG, Aldape KD, Yung WK, Salama SR, et al. Comprehensive, Integrative Genomic Analysis of Diffuse Lower-Grade Gliomas. The New England Journal of Medicine. 2015;372(26):2481–98.

9. Chen M, Sinha M, Luxon BA, Bresnick AR, O’Connor KL. Integrin alpha6beta4 controls the expression of genes associated with cell motility, invasion, and metastasis, including S100A4/metastasin. The Journal of biological chemistry. 2009;284(3):1484–94.

10. Chen S, Lewallen M, Xie T. Adhesion in the stem cell niche: biological roles and regulation. Development. 2013;140(2):255–65.

11. Cruz-Monserrate Z, Qiu S, Evers BM, O’Connor KL. Upregulation and redistribution of integrin alpha6beta4 expression occurs at an early stage in pancreatic adenocarcinoma progression. Mod Pathol. 2007;20(6):656–67.

12. Dang L, White DW, Gross S, Bennett BD, Bittinger MA, Driggers EM, et al. Cancer-associated IDH1 mutations produce 2-hydroxyglutarate. Nature. 2009;462(7274):739–44.

13. Dolecek TA, Propp JM, Stroup NE, Kruchko C. CBTRUS statistical report: primary brain and central nervous system tumors diagnosed in the United States in 2005-2009. Neurooncology. 2012;14 Suppl 5:v1–49.

14. Eckel-Passow JE, Lachance DH, Molinaro AM, Walsh KM, Decker PA, Sicotte H, et al. Glioma Groups Based on 1p/19q, IDH, and TERT Promoter Mutations in Tumors. The New England Journal of Medicine. 2015;372(26):2499–508.

15. Fischer I, Gagner JP, Law M, Newcomb EW, Zagzag D. Angiogenesis in gliomas: biology and molecular pathophysiology. Brain pathology. 2005;15(4):297–310.

16. Friedlander M, Brooks PC, Shaffer RW, Kincaid CM, Varner JA, Cheresh DA. Definition of two angiogenic pathways by distinct alpha v integrins. Science. 1995;270(5241):1500–2.

17. Gilbert MR, Liu Y, Neltner J, Pu H, Morris A, Sunkara M, et al. Autophagy and oxidative stress in gliomas with IDH1 mutations. Acta neuropathologica. 2013.

18. Guilhamon P, Eskandarpour M, Halai D, Wilson GA, Feber A, Teschendorff AE, et al. Meta-analysis of IDH-mutant cancers identifies EBF1 as an interaction partner for TET2. Nature communications. 2013;4:2166.

19. Guo W, Giancotti FG. Integrin signalling during tumour progression. Nature reviews Molecular cell biology. 2004;5(10):816–26.

20. Hale JS, Otvos B, Sinyuk M, Alvarado AG, Hitomi M, Stoltz K, et al. Cancer stem cellspecific scavenger receptor CD36 drives glioblastoma progression. Stem cells. 2014;32(7):1746–58.

21. Hartmann C, Meyer J, Balss J, Capper D, Mueller W, Christians A, et al. Type and frequency of IDH1 and IDH2 mutations are related to astrocytic and oligodendroglial differentiation and age: a study of 1,010 diffuse gliomas. Acta neuropathologica. 2009;118(4):469–74.

22. Hegi ME, Diserens AC, Gorlia T, Hamou MF, de Tribolet N, Weller M, et al. MGMT gene silencing and benefit from temozolomide in glioblastoma. The New England Journal of Medicine. 2005;352(10):997–1003.

23. Hitomi M, Deleyrolle LP, Mulkearns-Hubert EE, Jarrar A, Li M, Sinyuk M, et al. Differential connexin function enhances self-renewal in glioblastoma. Cell reports. 2015;11(7):1031–42.

24. Horbinski C. What do we know about IDH1/2 mutations so far, and how do we use it? Acta neuropathologica. 2013;125(5):621–36.

25. Hu Y, Smyth GK. ELDA: extreme limiting dilution analysis for comparing depleted and enriched populations in stem cell and other assays. Journal of immunological methods. 2009;347(1-2):70–8.

26. Ichimura K, Pearson DM, Kocialkowski S, Backlund LM, Chan R, Jones DT, et al. IDH1 mutations are present in the majority of common adult gliomas but rare in primary glioblastomas. Neuro-oncology. 2009;11(4):341–7.

27. Kurokawa A, Nagata M, Kitamura N, Noman AA, Ohnishi M, Ohyama T, et al. Diagnostic value of integrin alpha3, beta4, and beta5 gene expression levels for the clinical outcome of tongue squamous cell carcinoma. Cancer. 2008;112(6):1272–81.

28. Laffaire J, Everhard S, Idbaih A, Criniere E, Marie Y, de Reynies A, et al. Methylation profiling identifies 2 groups of gliomas according to their tumorigenesis. Neuro-oncology. 2011;13(1):84–98.

29. Lathia JD, Gallagher J, Heddleston JM, Wang J, Eyler CE, Macswords J, et al. Integrin alpha 6 regulates glioblastoma stem cells. Cell stem cell. 2010;6(5):421–32.

30. Lathia JD, Li M, Hall PE, Gallagher J, Hale JS, Wu Q, et al. Laminin alpha 2 enables glioblastoma stem cell growth. Ann Neurol. 2012;72(5):766–78.

31. Lathia JD, Li M, Sinyuk M, Alvarado AG, Flavahan WA, Stoltz K, et al. High-throughput flow cytometry screening reveals a role for junctional adhesion molecule a as a cancer stem cell maintenance factor. Cell reports. 2014;6(1):117–29.

32. Lathia JD, Mack SC, Mulkearns-Hubert EE, Valentim CL, Rich JN. Cancer stem cells in glioblastoma. Genes & development. 2015;29(12):1203–17.

33. Lathia JD, Patton B, Eckley DM, Magnus T, Mughal MR, Sasaki T, et al. Patterns of laminins and integrins in the embryonic ventricular zone of the CNS. The Journal of comparative neurology. 2007;505(6):630–43.

34. Laxton RC DL, Aizpurua M, Bodi I, King A. Co-deletion of 1p/19q is Strongly Correlated with a High Level of MGMT Promoter Methylation in High Grade Gliomas as Revealed by Pyrosequencing. J Mol Genet Med. 2015;9(1):1747–0862.

35. Lee SM, Park SY, Shin SW, Kil IS, Yang ES, Park JW. Silencing of cytosolic NADP(+)-dependent isocitrate dehydrogenase by small interfering RNA enhances the sensitivity of HeLa cells toward staurosporine. Free Radic Res. 2009;43(2):165–73.

36. Lipscomb EA, Mercurio AM. Mobilization and activation of a signaling competent alpha6beta4integrin underlies its contribution to carcinoma progression. Cancer metastasis reviews. 2005;24(3):413–23.

37. Lu C, Ward PS, Kapoor GS, Rohle D, Turcan S, Abdel-Wahab O, et al. IDH mutation impairs histone demethylation and results in a block to cell differentiation. Nature. 2012;483(7390):474–8.

38. Lu S, Simin K, Khan A, Mercurio AM. Analysis of integrin beta4 expression in human breast cancer: association with basal-like tumors and prognostic significance. Clinical cancer research. 2008;14(4):1050–8.

39. Mas-Moruno C, Rechenmacher F, Kessler H. Cilengitide: the first anti-angiogenic small molecule drug candidate design, synthesis and clinical evaluation. Anti-cancer agents in medicinal chemistry. 2010;10(10):753–68.

40. Mulholland S, Pearson DM, Hamoudi RA, Malley DS, Smith CM, Weaver JM, et al. MGMT CpG island is invariably methylated in adult astrocytic and oligodendroglial tumors with IDH1 or IDH2 mutations. International journal of cancer. 2012;131(5):1104–13.

41. Nikolopoulos SN, Blaikie P, Yoshioka T, Guo W, Giancotti FG. Integrin beta4 signaling promotes tumor angiogenesis. Cancer cell. 2004;6(5):471–83.

42. O’Connor KL, Chen M, Towers LN. Integrin alpha6beta4 cooperates with LPA signaling to stimulate Rac through AKAP-Lbc-mediated RhoA activation. American journal of physiology Cell physiology. 2012;302(3):C605–14.

43. O’Connor KL, Nguyen BK, Mercurio AM. RhoA function in lamellae formation and migration is regulated by the alpha6beta4 integrin and cAMP metabolism. The Journal of cell biology. 2000;148(2):253–8.

44. Otvos B, Silver DJ, Mulkearns-Hubert EE, Alvarado AG, Turaga SM, Sorensen MD, et al. Cancer Stem Cell-Secreted Macrophage Migration Inhibitory Factor Stimulates Myeloid Derived Suppressor Cell Function and Facilitates Glioblastoma Immune Evasion. Stem cells. 2016;34(8):2026–39.

45. Paulus W, Baur I, Schuppan D, Roggendorf W. Characterization of integrin receptors in normal and neoplastic human brain. Am J Pathol. 1993;143(1):154–63.

46. Pollack IF, Hamilton RL, Sobol RW, Nikiforova MN, Lyons-Weiler MA, LaFramboise WA, et al. IDH1 mutations are common in malignant gliomas arising in adolescents: a report from the Children’s Oncology Group. Child’s nervous system: ChNS: official journal of the International Society for Pediatric Neurosurgery. 2011;27(1):87–94.

47. Previtali S, Quattrini A, Nemni R, Truci G, Ducati A, Wrabetz L, et al. Alpha6 beta4 and alpha6 beta1 integrins in astrocytomas and other CNS tumors. Journal of neuropathology and experimental neurology. 1996;55(4):456–65.

48. Previtali SC, Quattrini A, Pardini CL, Nemni R, Feltri ML, Boncinelli E, et al. Laminin receptor alpha6beta4 integrin is highly expressed in ENU-induced glioma in rat. Glia. 1999;26(1):55–63.

49. Riemenschneider MJ, Mueller W, Betensky RA, Mohapatra G, Louis DN. In situ analysis of integrin and growth factor receptor signaling pathways in human glioblastomas suggests overlapping relationships with focal adhesion kinase activation. Am J Pathol. 2005;167(5):1379–87.

50. Sanson M, Marie Y, Paris S, Idbaih A, Laffaire J, Ducray F, et al. Isocitrate dehydrogenase 1 codon 132 mutation is an important prognostic biomarker in gliomas. Journal of clinical oncology. 2009;27(25):4150–4.

51. Schonberg DL, Miller TE, Wu Q, Flavahan WA, Das NK, Hale JS, et al. Preferential Iron Trafficking Characterizes Glioblastoma Stem-like Cells. Cancer cell. 2015;28(4):441–55.

52. Seguin L, Desgrosellier JS, Weis SM, Cheresh DA. Integrins and cancer: regulators of cancer stemness, metastasis, and drug resistance. Trends in cell biology. 2015;25(4):234–40.

53. Shaw LM, Rabinovitz I, Wang HH, Toker A, Mercurio AM. Activation of phosphoinositide 3-OH kinase by the alpha6beta4 integrin promotes carcinoma invasion. Cell. 1997;91(7):949–60.

54. Stewart RL, Carpenter BL, West DS, Knifley T, Liu L, Wang C, et al. S100A4 drives nonsmall cell lung cancer invasion, associates with poor prognosis, and is effectively targeted by the FDA-approved anti-helminthic agent niclosamide. Oncotarget. 2016;7(23):34630–42.

55. Stewart RL, O’Connor KL. Clinical significance of the integrin alpha6beta4 in human malignancies. Laboratory investigation; a journal of technical methods and pathology. 2015;95(9):976–86.

56. Stewart RL, West D, Wang C, Weiss HL, Gal T, Durbin EB, et al. Elevated integrin alpha6beta4 expression is associated with venous invasion and decreased overall survival in non-small cell lung cancer. Human pathology. 2016;54:174–83.

57. Stupp R, Mason WP, van den Bent MJ, Weller M, Fisher B, Taphoorn MJ, et al. Radiotherapy plus concomitant and adjuvant temozolomide for glioblastoma. The New England Journal of Medicine. 2005;352(10):987–96.

58. Takada Y, Ye X, Simon S. The integrins. Genome biology. 2007;8(5):215.

59. Tilghman J, Schiapparelli P, Lal B, Ying M, Quinones-Hinojosa A, Xia S, et al. Regulation of Glioblastoma Tumor-Propagating Cells by the Integrin Partner Tetraspanin CD151. Neoplasia. 2016;18(3):185–98.

60. Ward PS, Patel J, Wise DR, Abdel-Wahab O, Bennett BD, Coller HA, et al. The common feature of leukemia-associated IDH1 and IDH2 mutations is a neomorphic enzyme activity converting alpha-ketoglutarate to 2-hydroxyglutarate. Cancer cell. 2010;17(3):225–34.

61. Yan H, Parsons DW, Jin G, McLendon R, Rasheed BA, Yuan W, et al. IDH1 and IDH2 mutations in gliomas. The New England Journal of Medicine. 2009;360(8):765–73.

62. Zou P, Xu H, Chen P, Yan Q, Zhao L, Zhao P, et al. IDH1/IDH2 Mutations Define the Prognosis and Molecular Profiles of Patients with Gliomas: A Meta-Analysis. PloS one. 2013;8(7):e68782.

